# DNA damage in telophase leads to coalescence between segregated sister chromatid loci

**DOI:** 10.1101/437145

**Authors:** Jessel Ayra-Plasencia, Félix Machín

**Author notes:** Contact Unidad de Investigación, Hospital Universitario Nuestra Señora de Candelaria. Carretera del Rosario, 145. 38010. Santa Cruz de Tenerife, Spain.

## Abstract

The generation of DNA double strand breaks (DSBs) pose a high risk for the maintenance of the genome. Cells repair DSBs through two major mechanisms: non-homologous end joining (NHEJ) and homologous recombination (HR). HR is usually preferred when a sister chromatid is available, thus cells have coupled the activity of the cycling dependent kinase (CDK) to the selection of HR (Symington et al. 2014). Paradoxically, there is a window in the cell cycle where CDK is high despite a sister chromatid is not physically available for HR: late anaphase/telophase. We have here studied in budding yeast the response to DSBs generated in telophase by means of the radiomimetic drug phleomycin. We first show that phleomycin treatment activates the DNA damage response and leads to a delay in the telophase-to-G1 transition. Outstandingly, we also found a partial reversion of sister chromatid segregation, which includes approximation of spindle pole bodies (SPBs) and sister centromeres, *de novo* formation of anaphase bridges, trafficking of DNA back and forth through the cytokinetic plane and events of coalescence between segregated sister telomeres. We importantly show that phleomycin promotes a massive change in the structure and dynamic of mitotic microtubules (MTs), which coincides with dephosphorylation and re-localization of kinesin-5 Cin8. We propose that anaphase is not entirely irreversible and that there could still be a window to repair DSBs using the sister chromatid after segregation.

## Results & Discussion

We took advantage that *Saccharomyces cerevisiae* cells can be easily and stably arrested in telophase to check the DSB response at this cell cycle stage. We arrested cells in telophase using the broadly-used thermosensitive allele *cdc15-2*. Cdc15 is a key kinase in the Mitotic Exit Network (MEN) that allows cytokinesis and reduction of overall CDK activity, a hallmark of G1 (D’Amours & Amon 2004). We created randomly-distributed DSBs using phleomycin, a radiomimetic drug (Moore 1989). We first synchronously arrested in telophase *cdc15-2* strains that also carried YFP-labelled loci along the chromosome XII right arm (cXIIr; *tetO*/TetR-YFP system) (Quevedo et al. 2012). Treatment with phleomycin (10 μg/ml, 1h) caused hyperphosphorylation of Rad53 (Fig 1a), a classical marker for the activation of the DNA damage response (Finn et al. 2011). The degree of hyperphosphorylation was equivalent to those seen in G1- and G2/M-blocked cells, where Rad53 amplifies the corresponding checkpoint responses that delay G1-to-S transition and anaphase onset, respectively (Allen et al. 1994). We thus checked whether a telophase-to-G1 delay was also observed after phleomycin treatment. Indeed, concomitant removal of phleomycin and re-activation of Cdc15-2 (shift the temperature from 37°C to 25°C) showed a delay in both cytokinesis and G1 entry relative to cells which were not treated with phleomycin (mock treatment). For instance, plasma membrane ingression and resolution at the bud neck (abscission) was clear for ∼70% of cells just 1 h after Cdc15 re-activation (Fig 1b); note that most mother-daughter doublets remain together during a *cdc15-2* release, at least for the upcoming cell cycle (Quevedo et al. 2012). When telophase cells were treated with phleomycin, abscission was severally delayed; < 50% by 3h (Fig 1b). The telophase-to-G1 delay was also evident through three additional markers. Firstly, only G1 cells respond to the alpha-factor pheromone (αF) acquiring a schmoo-like morphology. We thus added αF after reactivating Cdc15 and found that less than 50% of phleomycin-treated cells had responded to αF by 3h, versus ∼75% of mock-treated cells (Fig 1c). Secondly, the CDK inhibitor Sic1 is only present in G1 cells (Schwob et al. 1994). We checked Sic1 levels after Cdc15 reactivation and found a clear delay in its production after phleomycin treatment (Fig 1d). Thirdly, flow cytometry (FACS) showed that the 2C content, expected in cells arrested in telophase, was long-lasting after phleomycin. By contrast, a mock-treated culture shortly turned this 2C peak into either 1C content (mother-daughter doublets are more easily separated during the rash treatment for FACS, provided that cytokinesis is completed) or 4C (DNA replication of the immediate progeny without mother-daughter separation) (Fig 1e).

**Figure 1.**
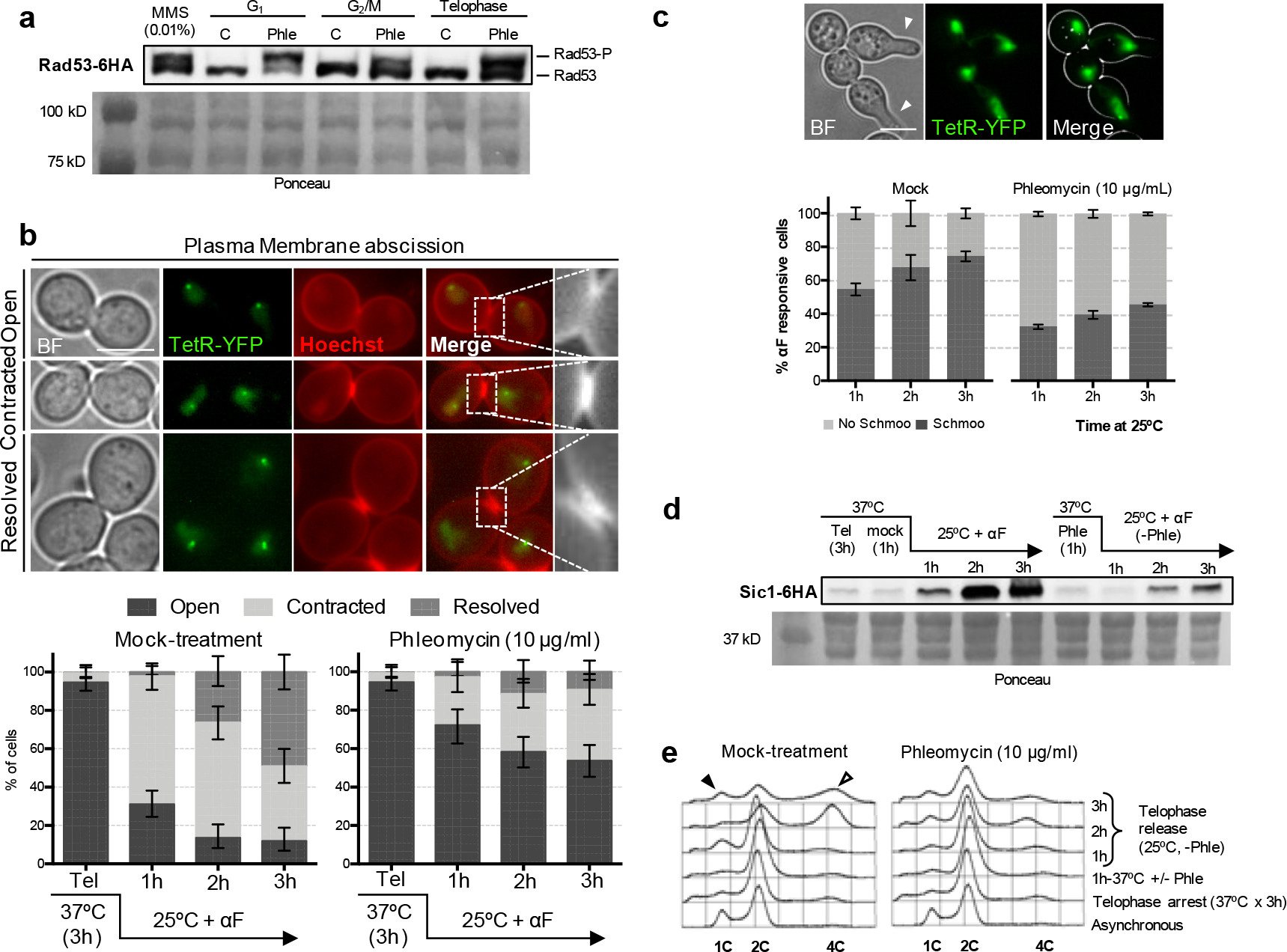
Phleomycin triggers a DNA Damage Response in telophase and a prolonged telophase-G1 delay. (**a**) Rad53 gets hyperphosphorylated in telophase after phleomycin treatment. Strain FM2329 was arrested for 3h at the indicated cell cycle stages before splitting the culture in two. Phleomycin (10 μg/mL) was added to one culture and cells were harvested after 1h for Western blot analysis. MMS (0.01%), added to an asynchronous culture, was used as Rad53 hyperphosphorylation control. C: Control (mock-treatment); Phle: Phleomycin-treated cells. The leftmost lane in the Ponceau staining corresponds to the protein weight markers. (**b**) Cytokinesis is delayed after DNA damage in telophase. Strain FM593 was arrested in telophase for 3 h, the culture split in two and one subculture treated with 10 μg/mL phleomycin. After 1 h, phleomycin was washed away and both cultures were released from the telophase block (shift to 25°C), taking samples for plasma membrane abscission analysis by using Hoechst staining every hour. The G1-blocking pheromone alpha-factor (αF) was added at the time of the telophase release to simplify cell outcomes. On the top, representative micrographs of telophase cells with different degrees of cytokinesis completion. At the bottom, charts showing the march of cytokinesis during the telophase release (± CI95). (**c**) Responsiveness to αF (as a marker of G1) is delayed after DNA damage in telophase. FM593 was treated as described in panel (b). On the top, a representative micrograph showing two telophase-like cells (binucleated dumbbells) in which one daughter cell responds to αF by acquiring the shmoo morphology. At the bottom, chart depicting the evolution of the αF-responsive cells after the telophase release (mean ± SEM, n=3). (**d**) Sic1 synthesis (G1 marker) is delayed after DNA damage in telophase. FM2323 was treated as described in panel (b), taking samples for Western blot analysis at the indicated time points. The leftmost lane in the Ponceau staining corresponds to the protein weight markers. (**e**) Cell separation and entry in a new S phase is blocked after DNA damage in telophase. FM593 was treated as described in panel (b). In this case though, αF addition was omitted. Samples were taken for FACS analysis at the indicated time points. DNA content (1C, 2C or 4C) is indicated under each FACS profile. Filled arrowhead points to the 1C peak coming from G1 daughter cells which have managed to complete cytokinesis and septation; hollow arrowhead points to the 4C peak, which comes from the cell subpopulation entering a new S phase before completing septation.

We next focussed on the behaviour of chromosome loci during and after phleomycin treatment in the telophase arrest. We started with cXII centromere, as a representative of the centromere cluster according to the Rabl configuration (Zimmer & Fabre 2011). We noted that, whereas mock-treated telophase cells maintained a constant distance of ∼8 μm between segregated sister centromeres, phleomycin caused a shortening of this distance to < 6 μm (Fig 2a). This approximation between sister centromeres occurred around 1h after phleomycin addition and was maintained for at least another hour upon phleomycin removal. Furthermore, we noted that the approximation was asymmetric (Fig 2b), and up to 25% of phleomycin-treated telophase cells had one sister centromere at the bud neck (versus < 5% in mock-treated cells; *p*<0.001, Fisher’s exact test). Similar shortening for the distance between sister loci was seen for the cXIIr telomere (Fig S2). Strikingly, we also observed coalescence between sister telomeres (Fig S2; ∼7% in phleomycin vs ∼2% in mock treatment; *p*<0.001, Fisher’s exact test), which was further confirmed through short-term videomicroscopy (Fig 2c; movies 1-3). Filming individual cells also showed acceleration of inter loci movement and how coalescence lasted longer than expected from simple Brownian motion (Fig 2c and d). Noteworthy, a general increase in loci movement in G2/M has been reported after DSB generation (Miné-Hattab & Rothstein 2012; Dion et al. 2012). Approximation and eventual coalescence of sister loci appeared to be general phenomena after DNA damage by phleomycin. Firstly, using the histone variant H2A2-mCherry, which labels all nuclear DNA, we confirmed that phleomycin treatment shortens the distance of the bulk of the segregated nuclear masses (Fig S3a). In addition, we observed confined trafficking of segregated DNA across the bud neck (Fig S3b and movies 4-6). This trafficking involved chromatin that appears partly depleted of histones (at least H2A) or is less condensed than the average segregated masses. Strikingly, phleomycin caused the formation of *de novo* histone-labelled anaphase bridges (Fig 2e). These bridges included chromatin confined in bulgy nuclear domains (Fig 2e, f and movie 7), as we described before in topoisomerase II (Top2) deficient cells arrested in telophase and in the *cdc14-1* block (Ramos-Pérez et al. 2017; Quevedo et al. 2012). Secondly, approximation and coalescence were also observed for the ribosomal DNA array (rDNA), coated with the rDNA binding protein Net1-eCFP (Fig 2g & S4). The rDNA is characterized by its repetitive nature and for nucleating the formation of the nucleolus, a membraneless intranuclear specialized organelle. Together, we conclude in this set of experiments that all segregated nuclear and nucleolar material gets closer after DNA damage. Under these circumstances, events of sister loci coalescence occur. We hypothesize that coalescence is favoured by both the approximation of the segregated genetic material and the acceleration of loci movement within the nucleus.

**Figure 2.**
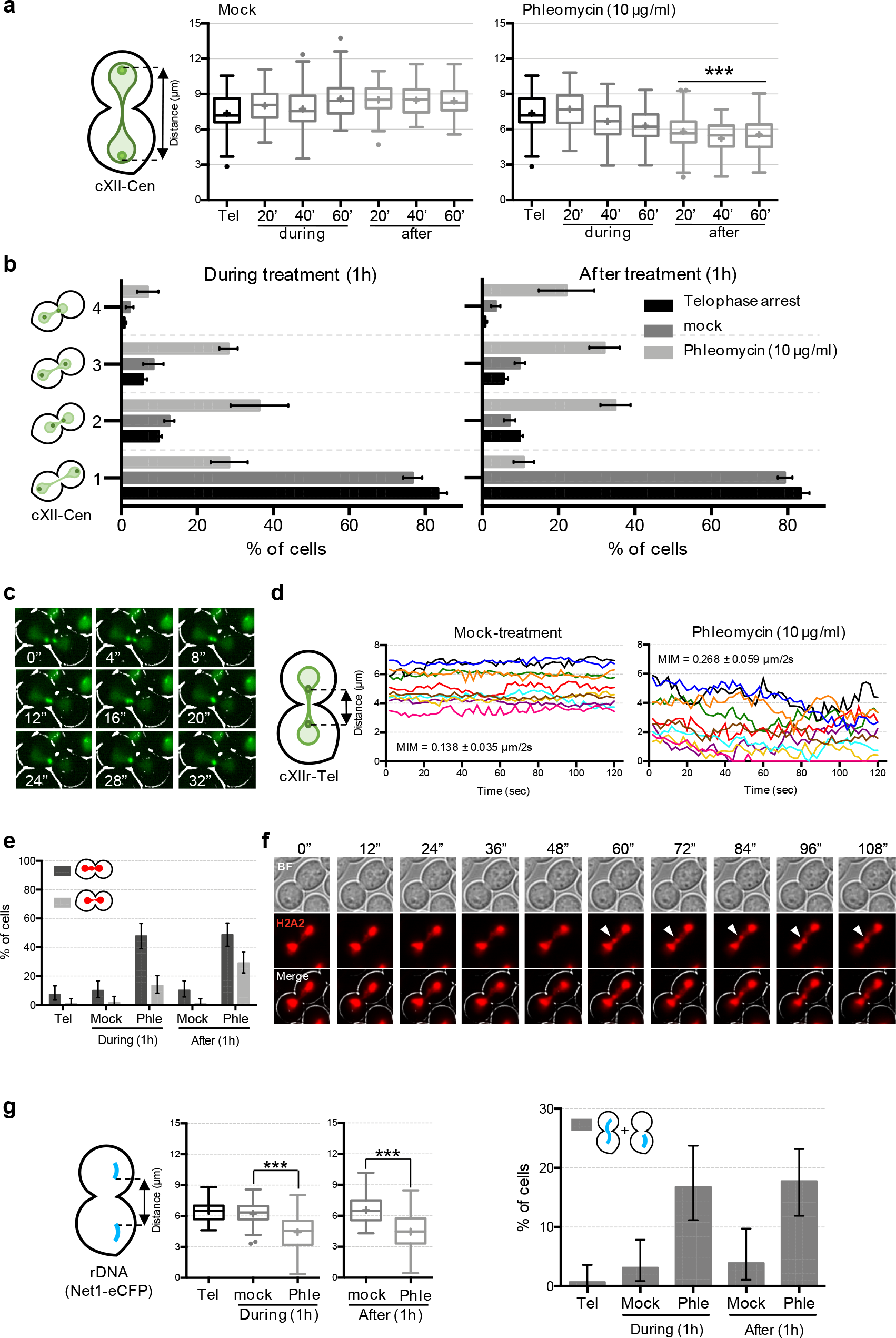
DNA damage in telophase leads to sister loci approximation, acceleration, coalescence and *de novo* formation of anaphase bridges. (**a**) Sister centromeres approach each other during and after DNA damage in telophase. FM593 was treated as in Fig 1e, except for the fact that the telophase block was maintained (37 °C) for 1 h after phleomycin was washed away. Samples were taken every 20’ during and after phleomycin (or mock) treatment and visualized under the microscope. Distance between sister cXII centromeres was measured and box-plotted for each time point (*** indicates p<0.0001 in mock/phleomycin comparisons at each time point; Mann-Whitney U Test). (**b**) Relative position of cXII sister centromeres from the former experiment was categorized as depicted on the left (mean ± SEM, n=3); category 1, both centromeres at near polar locations; category 2, closer centromeres with symmetrical distances to the bud neck; category 3, closer centromeres with asymmetrical distances to the bud neck; category 4, one sister centromere at the bud neck. (**c**) Sister telomeres can coalesce during DNA damage in telophase. FM588 was treated like in panel (a) and individual cells were filmed for 2 min after 1 h of phleomycin (or mock) addition. A representative cell where sister telomere coalescence was observed during the short movie. (**d**) Inter sister telomere movement accelerates after DNA damage. Kinetograms of 10 randomly selected FM588 cells. The mean interloci movement (MIM) during the 2 min movies is also displayed within the charts (mean ± SD, n=10 cells). The acceleration in interloci movement after phleomycin addition was statistically significant (p<0.0001, Student’s t test). (**e**) DNA damage in telophase brings about *de novo* chromatin bridges. FM2354 was treated like in (a) and morphology of the histone-labelled nuclear masses categorized in three: binuclear (not shown), with a gross/bulgy bridge (dark grey) and with a thinner/fainter histone-poor bridge (light grey). A representative experiment is shown (± CI95). (**f**) The *de novo* chromatin bridges are dynamics. FM2354 was filmed as in (c). A representative cell in which a bulgy bridge is dynamically formed from a histone-poor bridge (see also Fig S4). Filled arrowhead points to the bulge along the chromatin bridge. (**g**) Approximation and coalescence also occur for the rDNA/nucleolus. Strain FM2301 was treated like in (a). On the left, box-plots of minimum distances between sister rDNA signals under the indicated treatments (*** indicates p<0.0001; Mann-Whitney U Test). Single nucleolar signals were ignored for this calculation. On the right, bar chart for proportion of cells with a single nucleolus either in one cell body or stretched across the bud neck (± CI95).

Having observed the approximation of segregated sister loci, we next wondered about the cell forces underlying sister loci approximation. We consequently drove our attention to the spindle apparatus and engineered *cdc15-2* strains where we labelled the MTs (GFP-Tub1) and the SPBs (Spc42-mCherry), budding yeast equivalent to centrosomes. Phleomycin turned the elongated spindle, characteristic of telophase, into a rather dynamic star-like distribution (Fig 3a & movies 8-11). This new morphology points to a redistribution of Tubulin towards astral MTs, while nuclear MTs appears misaligned and with a weakened interpolar MT interaction. The change in the spindle morphology shortened the spindle length, which was confirmed by the approximation of the segregated SPBs from ∼9 to ∼6 μm (Fig 3b). The separation of SPBs in anaphase pulls attached centromeres apart, favouring the centromere-to-telomere segregation of sister chromatids (Machín et al. 2005). Co-visualization of SPBs and cXII sister centromeres showed that SPBs often headed centromeres in the approximation (Fig 3c), suggesting that either the strengthened astral microtubules or the weakened spindle indirectly drive sister loci approximation by pushing SPBs to each other. These results led us to check the behaviour of Cin8 upon DNA damage in telophase. Cin8 is a bidirectional mitotic kinesin-5 motor that slide away antiparallel interpolar microtubules, thus favouring spindle elongation in anaphase (Khmelinskii et al. 2009; Singh et al. 2018). Upon phleomycin treatment, Cin8 relocated from the spindle to two discrete foci in telophase-blocked cells (Fig 3d). These foci likely correspond to SPBs and/or kinetochore clusters (De Wulf et al. 2003; Shapira et al. 2017; Suzuki et al. 2018). Similar relocalization was observed in G2/M-blocked cells after phleomycin treatment (data not shown), suggesting a general Cin8 response to DSBs. Cin8 localization throughout the cell cycle depends on its phosphorylation status, with dephosphorylated Cin8 mostly locating at SPBs/kinetochores (Goldstein et al. 2017). We checked phosphorylation levels of Cin8 after DNA damage in telophase and found they are intermediate between S/G2 (fully dephosphorylated) and an unperturbed telophase (Fig 3e). Consequently, a partial dephosphorylation of Cin8 occurs upon DNA damage in telophase. We hypothesize that this Cin8 relocalization is likely a consequence of its novel minus-end-directed motility (Singh et al. 2018), and might reset the spindle to revert its elongation in cells already in late-anaphase.

**Figure 3.**
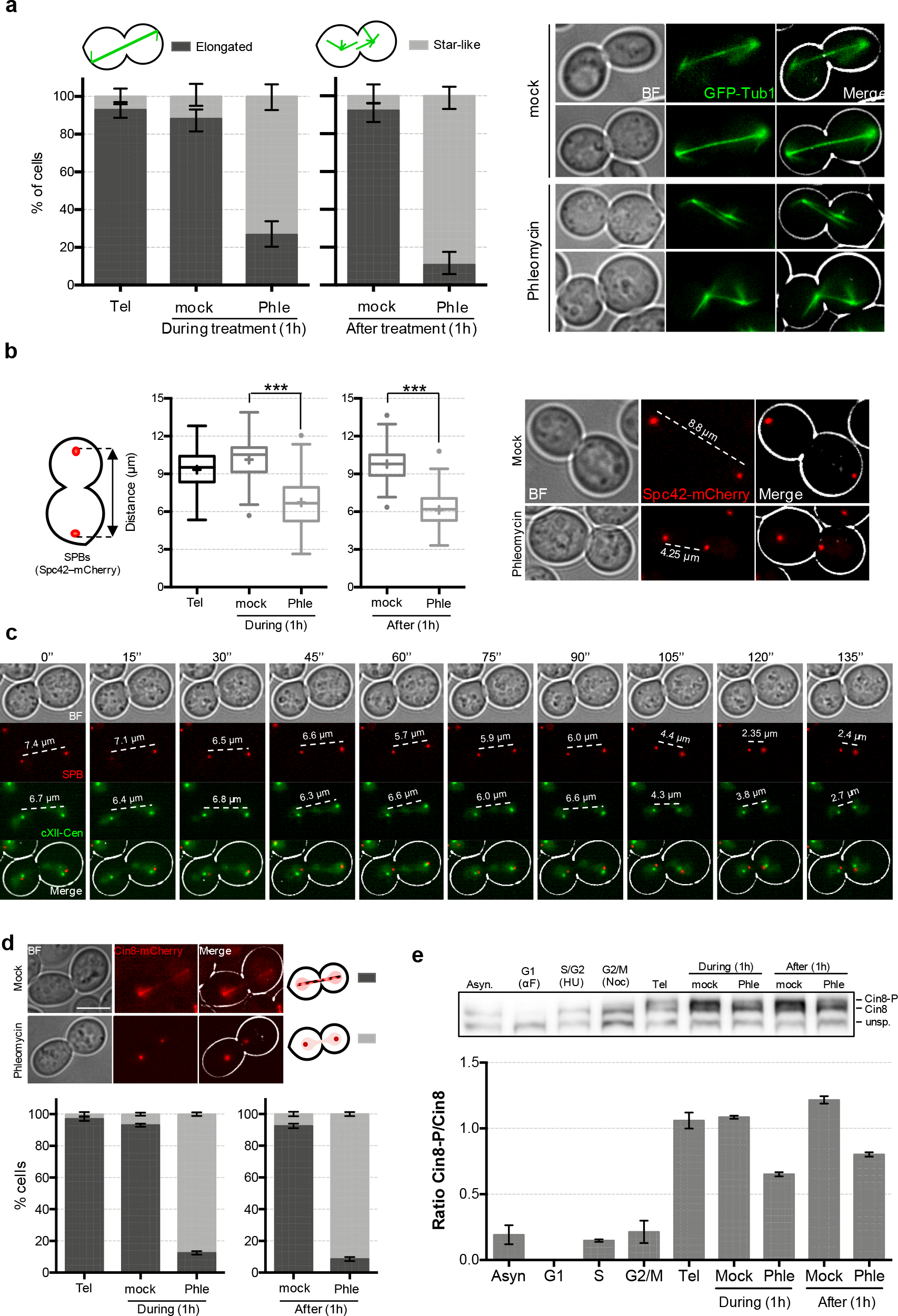
Dynamics of microtubules and kinesin-5 Cin8 after DNA damage in telophase. (**a**) Microtubules are repositioned after DNA damage in telophase. Strain FM2381 was treated as in Fig 2a. At the indicated conditions samples were taken and microtubules visualized under the microscope. Two categories were considered for quantification of the spindle: elongated and star-like; the latter being formed by multiple and shorter microtubule fibers arising from each SPB. (**b**) SPBs approach each other after DNA damage. Strain FM2316 was treated like in Fig 2a. On the left, box-plots of distances between SPBs under the indicated treatments (*** indicates p<0.0001; Mann-Whitney U Test). On the right, representative cells for the major phenotypes observed during each treatment. (**c**) SPBs often move ahead centromeres during the approximation that follows the DNA damage in telophase. Strain FM2316 was treated and filmed like in Fig 3c. (**d**) Kinesin-5 Cin8 relocalizes from the spindle into SPBs/kinetochores after DNA damage. Strain FM2317 was treated like in Fig 2a and checked under the microscope at indicated times. (**e**) A pool of Cin8 becomes dephoshorylated in telophase after DNA damage. Strain FM2335 was treated like in Fig 2a and, in addition, blocked at the indicated cell cycle stages. The upper picture shows Western blot of Cin8-myc, where myc signal appears as a triplet as reported before (Avunie-Masala et al. 2011). The lowest band is unspecific and serves as a loading control. The highest (slowest in PAGE migration) corresponds to the Cin8 phosphorylated forms (Cin8-P). Cin8 is absent in G1 (Avunie-Masala et al. 2011). The lower chart depicts quantification of the Cin8-P/Cin8 ratios (mean ± SEM, n=2).

Finally, we addressed whether the shift in MT distribution resulted in an active or passive mechanism for the approximation of the segregated material. We reasoned that Cin8 relocalization, together with enforced astral MTs, may result in new pulling forces to bring closer SPB/kinetochores/centromeres. Alternatively, weakening of interpolar MTs by sequestration of Cin8 out the spindle may passively allow approximation. To this aim, we studied the consequences of eliminating MTs in telophase-blocked cells, with or without concomitant DNA damage. Thus, nocodazole, a microtubule depolymerizing drug that does not exert DNA damage, mimicked most of the phenotypes just described for phleomycin; i.e., shortening of sister loci distances and acceleration of inter loci movement (Fig 4a, b & S5). In general, nocodazole masked the effect of phleomycin. For instance, nocodazole led to a more symmetric approximation of sister loci (compare categories 2 and 3 between Fig 2b and 4b), with double phleomycin-nocodazole treatment resembling the phenotype of just nocodazole. A similar relationship was seen for those cells where sister centromeres ended up within the same cell body (category 5 in Fig 4b). This was a very rare event in cells just treated with phleomycin, likely because astral MTs prevented SPBs from passing through the bud neck (movies 10 & 11). Importantly though, nocodazole and phleomycin had an additive effect on loci movement for a subset of cells (Fig S5b), demonstrating that other cell components aside from MTs participate in loci acceleration after the generation of DSBs in telophase. Noteworthy, phleomycin did not depolymerize microtubules. Firstly, the effect of nocodazole and phleomycin in telophase MTs was clearly different; nocodazole caused GFP-Tub1 to appear homogenously distributed throughout the cell, with signs of neither spindle nor astral microtubules (Fig S6a). Secondly, when added to an asynchronous culture, phleomycin arrested cells in G2/M with the characteristic metaphase GFP-Tub1 spindle (Fig S6b).

**Figure 4.**
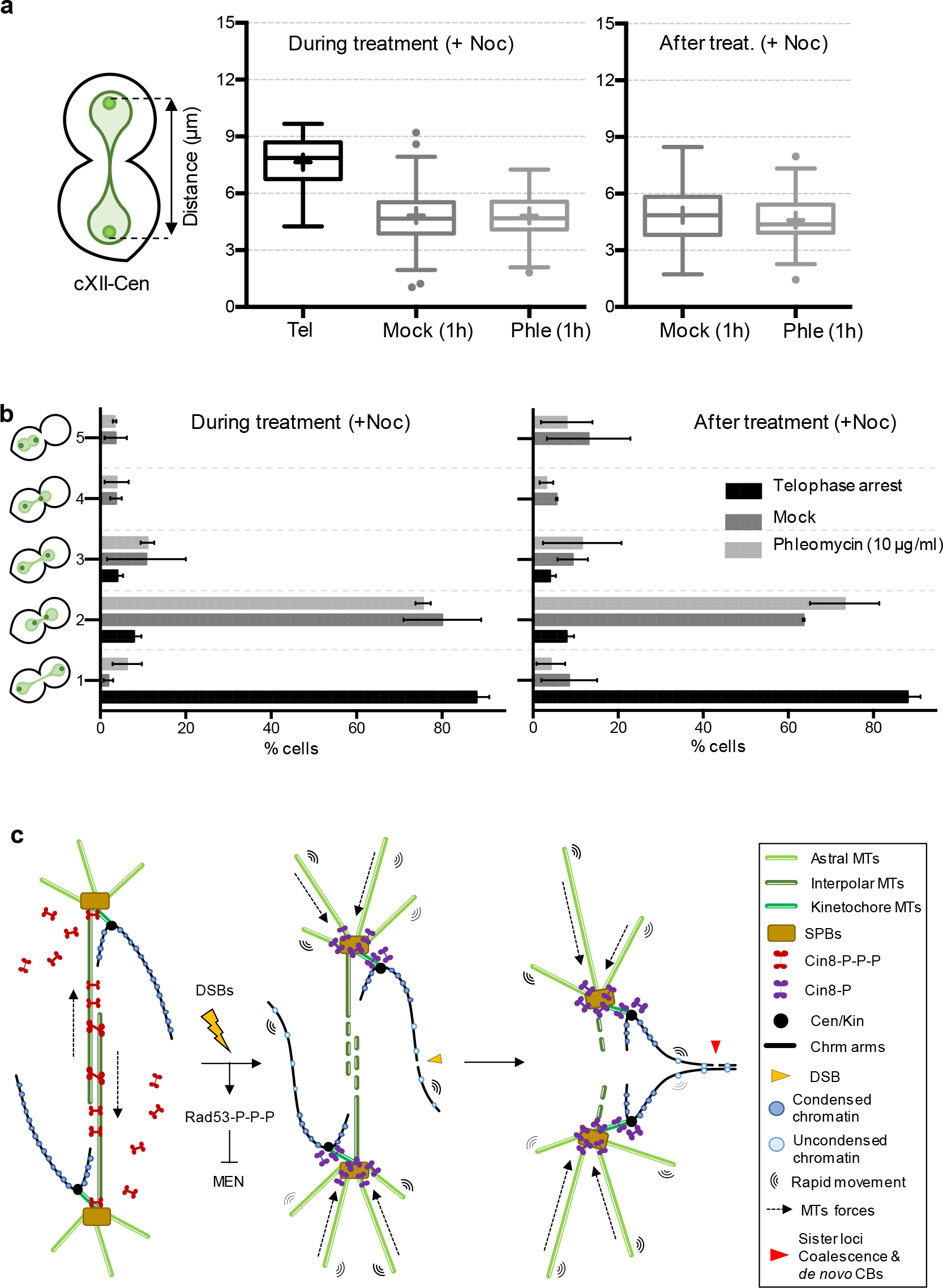
Microtubule depolymerization masks DNA damage effects on sister loci approximation in telophase. (**a**) Sister centromeres also approach each other after depolymerizing telophase microtubules, masking the effect of phleomycin. Strain FM593 was treated as in Fig 2a, except for the fact that nocodazole (Noc, 15 μg/mL) was added at the time of mock/phleomycin treatments and maintained after that. Distance between sister cXII centromeres was measured and box-plotted as in previous experiments. (**b**) Relative position of cXII sister centromeres from experiments like in panel (a) was categorized as indicated on the left (mean ± SEM, n=3). Categories 1 to 4 are described in Fig 2b. Category 5, two centromeres laying within the same cell body (very rare in Noc-free experiments). (**c**) Model for the effect of DNA damage on segregated sister chromatids. Segregated sister chromatids are maintained away from each other in telophase. Two complementary mechanisms aid to this aim. Firstly, an elongated spindle in maintained by the action of kinesin-5 Cin8 on interpolar microtubules (iMTs). Secondly, segregated sister chromatids are in a hypercondensed state (Machín et al. 2016). DNA damage (DSBs) locally mobilizes the affected chromatin through decondensation, so the histone-poor signal in *de novo* chromatin bridges, and globally accelerate loci movement. In addition, Cin8 is displaced out of iMTS by partial dephosphorylation, abrogating the spindle forces that keep SPBs far from each other. It is likely that enforced astral MTs also participate in bringing closer the SPBs. All these circumstances make possible for sister loci at chromosome arms to coalesce and form *de novo* chromatin bridges.

Altogether, we conclude that DSBs, at least those randomly generated through the DNA-cleaving agent phleomycin, can partially revert chromosome segregation in late anaphase. The results shown above question the irreversible nature of chromosome segregation, at least in budding yeast. Importantly, we provide mechanistic bases for this reversion (Fig 4c): (i) weakening of the elongated spindle, likely through dephosphorylation-dependent relocalization and sequestration of the bipolar kinesin-5 Cin8, which allows sister loci to get closer, and (ii) acceleration of loci movement, which increases the probability of closer sister loci to coalesce. We hypothesize that sister loci coalescence may provide a chance to repair DSBs through the efficient and error-free HR pathway. A window for such opportunity exists as we also demonstrate that DNA damage delays cytokinesis and telophase-G1 transition for more than 2 hours. Future work will be undertaken to validate this hypothesis.

## Materials and Methods

### Yeast strains and experimental procedures

All yeast strains used in this work are listed in Table S1. C-terminal tags were engineered using PCR methods (Smith & Burke 2014; Malcova et al. 2016). Strains were grown overnight in air orbital incubators at 25 °C in YEPD media (10 g/L yeast extract, 20 g/L peptone and 20 g/L glucose). To arrest cells in telophase, log-phase asynchronous cultures were adjusted to OD_600_ = 0.3 − 0.4 and the temperature shifted to 37 °C for 3 h. In most experiments, the arrested culture was split in two and one of them was treated with phleomycin (10 μg/mL) while the second was just treated with the vehicle (mock-treatment). After 1 h incubation, both cultures were washed twice with fresh YEPD and further incubated for 1-3 h to recover from DNA damage. In experiments aim to check the telophase-G1 transition, temperature was shifted back to 25 °C to allow Cdc15-2 re-activation. To simplify morphological outcomes during the *cdc15-2* release, the alpha factor pheromone (αF) was added after the washing steps unless stated otherwise (50 ng/mL; all strains are *bar1*, so hypersensitive to αF). For experiments other than telophase-G1 time courses, telophase arrest was maintained after phleomycin/mock treatments by keeping the temperature at 37 °C. Particular experiments such as plasma membrane ingression and responsiveness to αF have been described before (Quevedo et al. 2012; Ramos-Pérez et al. 2017). To arrest cells in G1, 50 ng/mL αF were directly added to an asynchronous culture growing at 25 °C and incubated at that temperature for 3 h. To arrest cells in G2/M, 15 μg/mL Nocodazole was added instead of αF. To arrest cells in S/G2, either 0.1% v/v methyl methanesulfonate (MMS) or 0.2 M hydroxyurea (HU) were added instead.

### Microscopy

A fully-motorized Leica DMI6000B wide-field fluorescence microscope was used in all experiments. In time courses, a stack of 20 z-focal plane images (0.3 μm depth) were collected using a 63×/1.30 immersion objective and an ultrasensitive DFC 350 digital camera. Micrographs were taken from freshly collected cells without further processing; 200-300 cells were quantified per experimental data point. Video-microscopy was also performed in freshly collected cells in a single focal plane (no more than 2 minutes, time frames of 2 seconds). The AF6000 (Leica) and Fiji (NIH) softwares were used for image processing and quantifications. The distances between sister loci and SPBs, as well as minimum distances between segregated rDNA and histone-labelled nuclear masses, were measured manually with the AF6000 software. Mean interloci movement (MIM) was calculated from the cumulative absolute variation of distances during videomicroscopy recording divided by the number of frames: (Σ|d_f_ - d_f−1_|)/n; where d is distance, f is the frame number, n is total number of frames, and the summation goes from f=2 to f=n. Coalescent events were not considered for calculations.

### Western blots and flow cytometry

For western blotting, 10 ml of the yeast liquid culture were collected to extract total protein using the trichloroacetic acid (TCA) method. Briefly, cell pellets were fixed in 2 mL of 20% TCA. After centrifugation (2,500 *g* for 3 min), cells were resuspended in 100 μL 20% TCA and ∼200 mg of glass beads were added. After 3 minutes of breakage by vortex, extra 200 μL 5% TCA were added to the mix. Samples were then centrifuged (2,500 *g* for 5 min) and pellets were resuspended in 100 μL of PAGE Laemmli Sample buffer mixed with 50 μL TE 1X pH 8.0. Finally, tubes were boiled for 3 min at 95 °C and pelleted again, being subjected to quantification with a Qubit 4 Fluorometer (Thermo Fisher Scientific, Q33227). Proteins were resolved in 10% (7.5% for Rad53 hyperphosphorylation assay) SDS-PAGE gels and transferred to PVFD membranes. The HA epitope was recognized with a primary mouse monoclonal anti-HA antibody (Sigma-Aldrich, H9658; 1:5,000); and the myc epitope was recognized with a primary mouse monoclonal anti-myc antibody (Sigma-Aldrich, M4439; 1:5,000). A polyclonal goat anti-mouse conjugated to alkaline phosphatase (Promega, S3721; 1:10,000) was used as secondary antibody.

Chemiluminescence method was selected to detection, using the CDP-Star reagent (GE Healthcare, RPN3682) and a Vilber-Lourmat Fusion Solo S documentation chamber. The membrane was finally stained with Ponceau S-solution for a loading reference. Flow cytometry to determine DNA content was performed as described before (García-Luis & Machín 2014).

### Data representation and statistics

Bar charts represent proportions of cells which have been categorized (e.g., relative position of sister loci, plasma membrane ingression, etc). Error bars in these charts generally depict the standard error of the mean (SEM), with the aim of quickly showing the interexperimental variability. At least 3 experiments, performed in different days, were considered for each figure panel. In case only one representative experiment is shown, error bars represent exact 95% confidence interval (CI95) of the proportion. Continuous data (e.g., interloci distance in μm) were represented in box-plots; the center line depicts the medians, the cross depicts the mean, box limits indicate the 25^th^ and 75^th^ percentile, whiskers extend to the 5^th^ and 95^th^, and dots represent outliers.

R software (https://www.r-project.org/) was used for statistical tests. Differences between experimental data points with continuous data were estimated through a Mann-Whitney U Test. Differences between experimental data points with categorical data were estimated through a Fisher’s exact test. In this case, cells counted in all three independent experiments were pooled (>500 cells per data point) to make the contingency tables. All reported *p* values are two-tailed.

## Acknowledgements

This work was supported by Spanish Ministry of Economy, Industry and Competitiveness (research grants BFU2015-63902-R and BFU2017-83954-R to FM). Both grants were co-financed with the European Commission’s ERDF structural funds.

## Author contributions

F.M. conceived the original project. J.A-P. performed all the experimental work and prepared the figures. F.M. and J.A-P. planned and analysed the experiments. F.M. wrote the paper.

